# Molecular Characterisation of the Synovial Fluid Microbiome in Rheumatoid Arthritis Patients and Healthy Control Subjects

**DOI:** 10.1101/405613

**Authors:** Dargham Bayan Mohsen Hammad, Veranja Liyanapathirana, Daniel Paul Tonge

## Abstract

The colonisation of specific body sites in contact with the external environment by microorganisms is both well-described and universally accepted, whereas, the existence of microbial evidence in other “classically sterile” locations including the blood, synovial space, and lungs, is a relatively new concept. Increasingly, a role for the microbiome in disease is being considered, and it is therefore necessary to increase our understanding of these. To date, little data support the existence of a “synovial fluid microbiome”.

**Methods:** The presence and identity of bacterial and fungal DNA in the synovial fluid of rheumatoid arthritis (RA) patients and healthy control subjects was investigated through amplification and sequencing of the bacterial 16S rRNA gene and fungal internal transcribed spacer region 2 respectively. Synovial fluid concentrations of the cytokines IL-6, IL-17A, IL22 and IL-23 were determined by ELISA.

**Results:** Bacterial 16S rRNA genes were detected in 87.5% RA patients, and all healthy control subjects. At the phylum level, the microbiome was predominated by *Proteobacteria* (Control = 83.5%, RA = 79.3%) and *Firmicutes* (Control = 16.1%, RA = 20.3%), and to a much lesser extent, *Actinobacteria* (Control = 0.2%, RA = 0.3%) and *Bacteroidetes* (Control = 0.1%, RA = 0.1%). Fungal DNA was identified in 75% RA samples, and 88.8% healthy controls. At the phylum level, synovial fluid was predominated by members of the Basidiomycota (Control = 53.9%, RA = 46.9%) and Ascomycota (Control = 35.1%, RA = 50.8%) phyla. Statistical analysis revealed key taxa that were differentially present or abundant dependent on disease status.

**Conclusions:** This study reports the presence of a synovial fluid microbiome, and determines that this is modulated by disease status (RA) as are other classical microbiome niches.

## Introduction

### The Microbiome

The term “microbiome” describes the entire habitat, including the microorganisms (bacteria, archaea, lower and higher eukaryotes, and viruses), their genomes, and the surrounding environmental conditions. Herein, we utilise this term to denote the existence of a microbial community evidenced through the survey of 16S rRNA genes. In contrast, the term “microbiota” refers to the assemblage of viable microorganisms that comprise these communities [1]. Whilst previous estimates stated that the number of microbial cells present in our microbiota exceeded our own by approximately one order of magnitude, current estimates are more conservative and suggest that the number of bacterial cells in the human body is roughly equal to the number of human cells, representing a mass of approximately 0.2 kg [2]. In either case, the microbiota represents a significant source of non-host biological material. The microbiota undertakes essential biological processes and thus it is unsurprising that a number of disease states are associated with changes in microbiome composition, termed “dysbiosis”. Whilst the colonisation of specific body sites in contact with the external environment (such as the gastrointestinal tract, skin and vagina) by microorganisms is both well-described and universally accepted [3], the existence of microorganisms and or microbial DNA in other “classically sterile” locations including the blood [4], synovial space [5], and lungs [6–9], is a relatively new concept.

### Rheumatoid Arthritis and Dysbiosis

Rheumatoid arthritis (RA) is a chronic autoimmune disease that causes synovial joint inflammation leading to significant joint pain, swelling, and significant disability with increased morbidity and mortality. RA is estimated to affect 0.5 - 1% of the population worldwide [10]. RA aetiology still remains unclear. Although the autoimmune hypothesis is well established the causes generating this self-directed immune reaction are still poorly understood. Although convention still suggests otherwise, an overwhelming volume of epidemiological and experimental data indicates a microbial origin for RA (reviewed extensively by Pretorius and colleagues [11]) and therefore increasingly, a role for the altered microbiome composition in disease initiation and progression is considered. To date, RA has been associated with dysbiosis of the oral [12, 13], gastrointestinal [12, 14–22], and respiratory [23–25] microbiomes and associated with the presence of specific organisms (most commonly *P. mirabilis*) within the urinary tract [26–29], suggesting that RA results in (or from) pan-microbiome dysbiosis. Studies have further begun to associate RA, or stages thereof, with the presence or absence of specific bacteria, including the expansion of rare bacterial linages [16]. Such findings have the potential to increase our disease understanding, and further suggest that the microbiome may afford a valuable source of novel biomarkers and or novel targets for therapeutic modulation.

The classical assertion that RA lacks a microbial component stems from the generalised inability of clinicians and scientists to observe microbial colonies on synthetic media when samples from RA patients are subjected to routine culture. Such assessments crudely characterise organisms as “*alive*” or “*dead*” (or perhaps more accurately, present or not present) based upon this colony forming ability, yet do not routinely recognise the potential for the presence of viable-non culturable bacteria. Furthermore, extensive evidence now provides for the existence of bacterial DNA and RNA in classically sterile areas leaving the possibility of; (a) microbial translocation from classical niches (e.g. the gastrointestinal tract) followed by opsonisation leaving only the nucleic acid, (b) the passage of cell-free microbial nucleic acid through the circulatory system, (c) microbial translocation followed by a state of dormancy due to unfavourable environmental conditions. With relevance to the first two scenarios, it is well-established that the immune system is able to differentiate self from non-self-nucleic acid (DNA and RNA) through specific pattern recognition receptions [30], and that foreign DNA is recognised by TLR9, and RNA is recognised by TLR3 on immune cells, resulting in the upregulation of type I interferons and various pro-inflammatory cytokines [31] (also associated with rheumatoid arthritis) [32]. TLR9 is found in the endosome of peripheral blood mononuclear cells, such as monocytes, macrophages, T, B, and NK cells, whereas TLR3 is expressed only in the endosomes of myeloid derived cells such as dendritic cells and monocytes [33]. With relevance to the latter scenario, dormant or VBNC bacterial cells may persist within the joints of arthritic patients, undetected by routine culture, whilst retaining the ability to shed inflammatory agents such as lipopolysaccharide (LPS) and other antigenic components [11], yet such samples may present as culture-negative. Indeed, bacterial LPS is able to simulate many of the common RA-associated cytokines [34].

### Evidence of Bacteria in the Synovial Fluid: Origin and Transport

Under normal physiological conditions, one expects the synovial space to be sterile. Indeed, the presence of viable bacteria results in a diagnosis of septic arthritis, a condition regarded as a true medical emergency [35]. Nevertheless, numerous studies report the presence of microorganisms, or microorganism-derived genetic material in the synovial fluid of RA patients and healthy control subjects [36–42]. Interestingly, these studies overwhelmingly identify bacteria most commonly found to reside within the oral cavity, and these findings are accompanied by evidence demonstrating the presence of these DNAs and or antibodies directed against the originating organism in the blood [36, 37, 43–46]. On the basis of these findings, we suggest that bacteria / bacterial nucleic acid may reach the synovial space via the blood (as evidenced by their concurrent presence in both fluids), and further, that the inflammatory environment of the synovial tissues may facilitate the trapping of these bacterial DNAs, increasing their apparent concentration in this location [37].

Despite extensive epidemiological and experimental evidence supporting a microbial cause in RA, mounting evidence linking infection of the oral tissues with RA [23, 47–58], and data demonstrating the appearance of microbial DNA from distant microbiome niches in the synovial space (see above), there has been limited study of the bacterial community in this compartment [5] and no global characterisation of the fungal DNA present in this location to date. To this end, we investigated the presence of bacterial (via sequencing of the 16S rRNA gene) and fungal (via sequencing of the ITS2 region) DNA, in the synovial fluid of human RA patients and healthy control subjects using cutting-edge next generation sequencing and bioinformatic techniques. Furthermore, we evaluated any apparent changes in the bacterial or fungal communities detected, in the context of key markers of synovial inflammation (IL-6, IL-17A, IL-22 and IL-23).

## Materials and Methods

### Donor Population

This study investigated the presence of bacterial and fungal DNA in synovial fluid samples from sixteen rheumatoid arthritis patients, and nine sex and BMI-matched healthy control (disease-free) individuals who donated a sample of their synovial fluid for research purposes. Synovial fluid samples were aspirated from the knee joint using aseptic technique, transferred to a sterile micro-centrifuge tube, and stored at −80°C prior to further analysis. Samples were procured from Sera Laboratories Limited.

The Independent Investigational Review Board Inc. (BRI-0722) ethically approved sample collection by Sera Laboratories Limited from human donors giving informed written consent. Furthermore, the authors obtained ethical approval from Keele University Ethical Review Panel 3 for the study reported herein. All experiments were performed in accordance with relevant guidelines and regulations.

### Measuring synovial fluid cytokines

Markers of inflammation were measured in synovial fluid utilising the Human Magnetic Luminex Screening Assay following the manufacturer’s instructions (R&D Systems, Minneapolis, USA). Levels of IL-6, IL-17A, IL-22, and IL-23 were analysed by Luminex kit LXSAHM-04. Prior to analysis, synovial fluid samples were centrifuged at 16,000 x g for four minutes and diluted in a 1:2 ratio by adding 25 μl of sample to 25 μl of Assay Buffer prior to analysis.

### Microbiome Characterisation

Amplification and sequencing of the bacterial 16S rRNA variable region 4 and fungal ITS region was used to characterise the bacterial and fungal DNA present in the synovial fluid samples.

The bacterial 16S rRNA and fungal ITS2 genes were amplified by direct PCR utilising the oligonucleotide primers detailed in (**Table 1**). Such approaches lack the need for a separate DNA extraction phase, often associated with the introduction of contaminants that impact upon all downstream applications. A first round of PCR using oligonucleotide primers 16SV4 F/R and ITS2 F/R was carried out with four microliters of each synovial fluid sample as template in a final volume of 20µl. Reactions comprised 10µl Phusion blood PCR buffer (Thermofisher), 0.4µl (2 U) Phusion blood DNA polymerase, 1µl of each primer (10μM), and 3.6µl of nuclease-free water that had been subject to 15 minutes UV-irradiation. A negative control reaction in which synovial fluid was substituted with UV-irradiated nuclease-free water was run in parallel with our experimental samples to confirm that none of the reagents were contaminated by target DNA. Cycling parameters comprised an initial denaturation step performed at 98°C for 5 minutes, followed by 33 cycles of denaturation (98°C, 10 seconds), annealing (55°C, 5 seconds) and extension (72°C, 15 seconds). A final elongation of 7 minutes at 72°C completed the reaction. Following gel electrophoresis of 5μl of each resulting PCR product to confirm successful amplification, the remainder (15μl) was purified of excess primer and PCR reagents using the QIAquick PCR Purification kit.

**Table 1:**
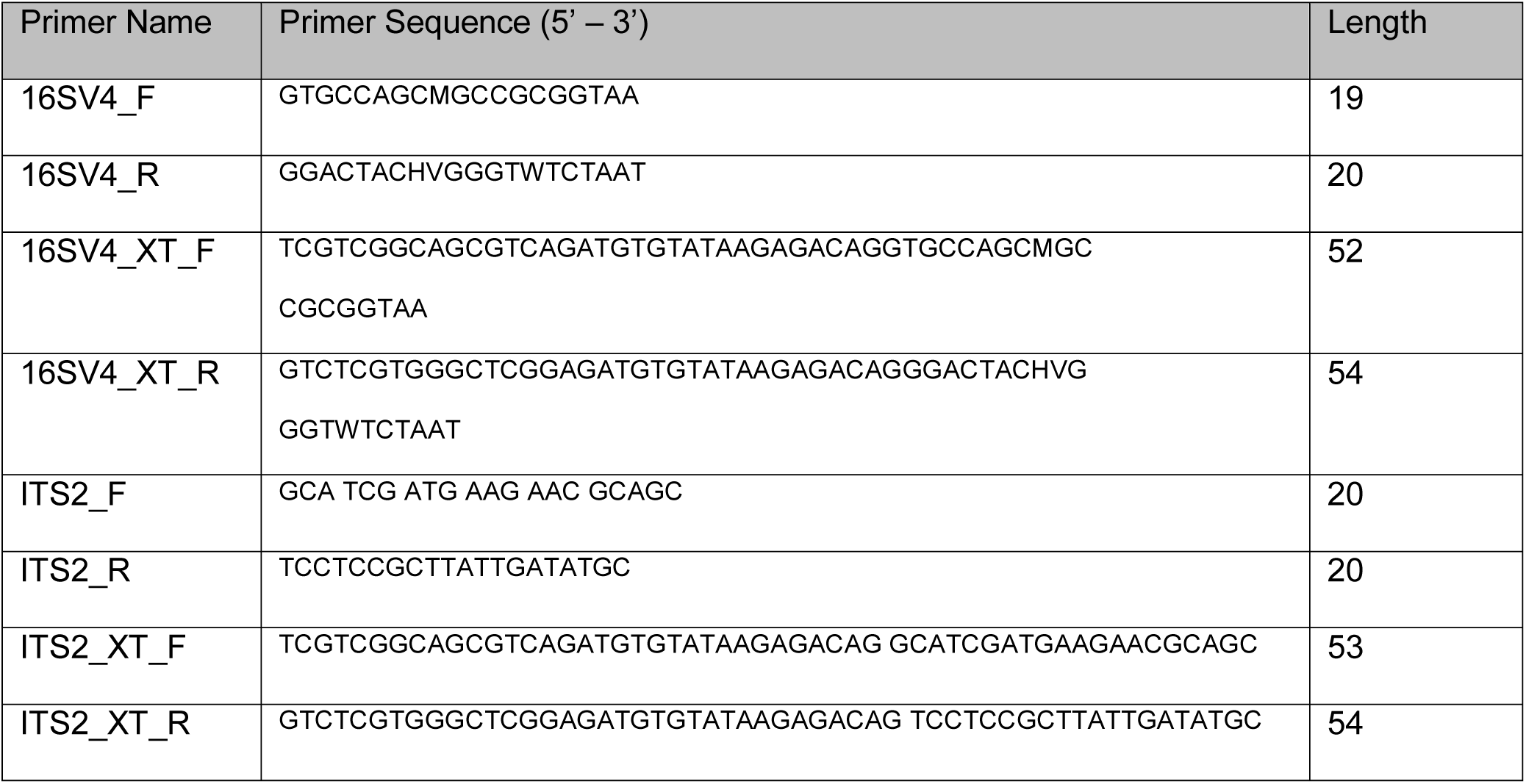
Oligonucleotide primers used in this study

In order to add sequencing adapters to facilitate MiSeq library preparation, a second round of PCR amplification was conducted in a total volume of 50 μl comprising 10µl 5X Platinum Super-Fi Buffer, 1µl 10mM dNTP mixture, 0.5µl Platinum Super-Fi polymerase, 2.5µl of each XT_tagged primer (16SV4_XT F/R or ITS2_XT F/R), 5µL from the successful first round PCR reaction and 38.5 µL of UV-irradiated nuclease-free water. Cycling parameters comprised an initial denaturation step performed at 98°C for 2 minutes, followed by 7 cycles of denaturation (98°C, 10 seconds), and annealing / extension (72°C, 20 seconds) extension (72°C, 15 seconds). A final elongation of 5 minutes at 72°C completed the reaction. Two controls were run in parallel with this process and these comprised a standard PCR negative control (in which template was replaced with UV irradiated molecular biology grade water) and the first round PCR control which was progressed alongside our experimental samples. All PCR products and our two control reactions were purified using AMPure XP magnetic beads (Agencourt) at a ratio of 0.8 beads to sample (v/v), eluted in 20 μl of UV-irradiated nuclease-free water, and visualised by agarose gel electrophoresis and ehidium bromide staining as above. All PCR negative reactions were subject to high sensitivity DNA quantification using the Qubit 3.0 hsDNA kit (Invitrogen) to quantitate any DNA in these samples.

Amplicons were barcoded using the Nextera DNA library kit, multiplexed for efficiency, and sequenced using the Ilumina MiSeq system with a 250bp paired-end read metric. Bioinformatic analysis was performed using QIIME implemented as part of the Nephele 16S / ITS paired-end QIIME pipeline using open reference clustering against the SILVA database for bacteria and the ITS database for fungi at a sequence identity of 99%. All other parameters remained as default. The statistical significance of differences in the abundance of individual bacteria and fungi between the RA and control donors was determined by applying a two-tailed, Mann Whitney test using GraphPad Prism V8.0.

## Results

### Clinical features of donors and results of 16S rRNA and ITS2 PCR amplification

Synovial fluid samples were obtained from 25 human donors. Sixteen patients were diagnosed with RA, and of these, 8 were male and 8 were female. The RA patients’ ages ranged from 52 to 74 years, with a mean of 65.3 years. Nine control synovial fluid samples were obtained from 5 males and 4 females. Their ages ranged from 50 to 68 years, with a mean of 61.0 years. There were no significant differences in the ages of the two cohorts (Unpaired T-test; P = > 0.05) (**Table 2**).

**Table 2:** Characteristics of the patient population.

| Patient ID# | Age | Diagnosis | 16S rRNA<br>PCR | ITS2<br>PCR |
| --- | --- | --- | --- | --- |
| 1339 | 61 - 65 | RA | + | - |
| 1340 | 66 - 70 | RA | + | + |
| 1341 | 66 - 70 | RA | + | + |
| 1342 | 66 - 70 | RA | + | - |
| 1343 | 66 - 70 | RA | + | - |
| 1344 | 66 - 70 | RA | + | - |
| 1345 | 56 - 60 | RA | + | + |
| 1346 | 50 - 55 | RA | + | + |
| 1347 | 50 - 55 | RA | + | + |
| 1348 | 66 - 70 | RA | + | + |
| 1349 | 71 - 75 | RA | + | + |
| 1350 | 66 - 70 | RA | + | + |
| BRH1095336 | 66 - 70 | RA | + | + |
| BRH1095337 | 66 - 70 | RA | - | + |
| BRH1095338 | 66 - 70 | RA | - | + |
| BRH1095339 | 66 - 70 | RA | + | + |
| 1351 | 56 - 60 | Healthy | + | + |
| 1352 | 61 - 65 | Healthy | + | - |
| 1353 | 61 - 65 | Healthy | + | + |
| 1354 | 50 - 55 | Healthy | + | + |
| 1355 | 50 - 55 | Healthy | + | + |
| 1356 | 71 - 75 | Healthy | + | + |
| 1357 | 61 - 65 | Healthy | + | + |
| 1358 | 50 - 55 | Healthy | + | + |
| 1359 | 66 - 70 | Healthy | + | + |

Utilising PCR amplification, bacterial 16S DNA was detected in the synovial fluid of 14 of out 16 patients with RA (87.5%), and in 9 out of 9 (100%) healthy control subjects. ITS2 amplification, indicative of the presence of fungi, was detected in 12 of 16 (75%) RA samples, and 8 out of 9 (89%) healthy control samples. Our various experimental controls designed to identify target contamination amongst the molecular reagents and plasticware used consistently failed to produce a visible band following PCR and agarose gel electrophoresis. Furthermore, these samples were subjected to DNA quantification using the Qubit 3.0 high-sensitivity DNA kit (Invitrogen) and were found to be below the limit of detection in all cases.

### Assessment of inflammatory markers in synovial fluid

Synovial fluid concentrations of interleukin 6 (IL-6), 17A (IL-17A), 22 (IL-22) and 23 (IL-23) were measured using the Luminex system as described earlier. Mean interleukin levels in healthy control and RA patient synovial fluid were significantly different, with cytokines being present at higher levels in the RA synovial fluid in all cases (Unpaired T-test) (**Table 3**).

**Table 3.** Differences in cytokine levels among

| Cytokine | Control mean (SD) (pg/mL) | RA mean (SD) (pg/mL) | P value (pg/mL) |
| --- | --- | --- | --- |
| IL-6 | 156.43 (100.30) | 1257.80 (905.66) | <0.005 |
| IL-17A | 11.68 (3.92) | 65.44 (16.51) | <0.005 |
| IL-22 | 220.14 (85.02) | 873.00 (376.46) | <0.005 |
| IL-23 | 145.64 (64.43) | 296.21 (97.07) | <0.005 |

### Characterisation of Bacterial populations via 16S rDNA sequencing of synovial fluid

The presence of bacterial DNA in synovial fluid was evaluated via PCR amplification and sequencing of the bacterial 16S rRNA gene, variable region 4. An average of 75,000 reads were generated for each of the samples; 70,658 reads in the RA samples, and 80,533 reads in the control samples. Although the control samples generated more reads on average, this difference was not statistically significant (Unpaired T test; P = > 0.05), and nevertheless, rarefaction was used prior to differential abundance analysis to control for differences in sequencing depth. Following taxonomic classification using the Nephele platform as described above, our first analytical approach utilised Principal Coordinates Analysis (PCoA) to reduce the complexity of the data obtained, and to visualise any obvious separation between the various experimental samples (**Figure 1**).

**Figure 1.**
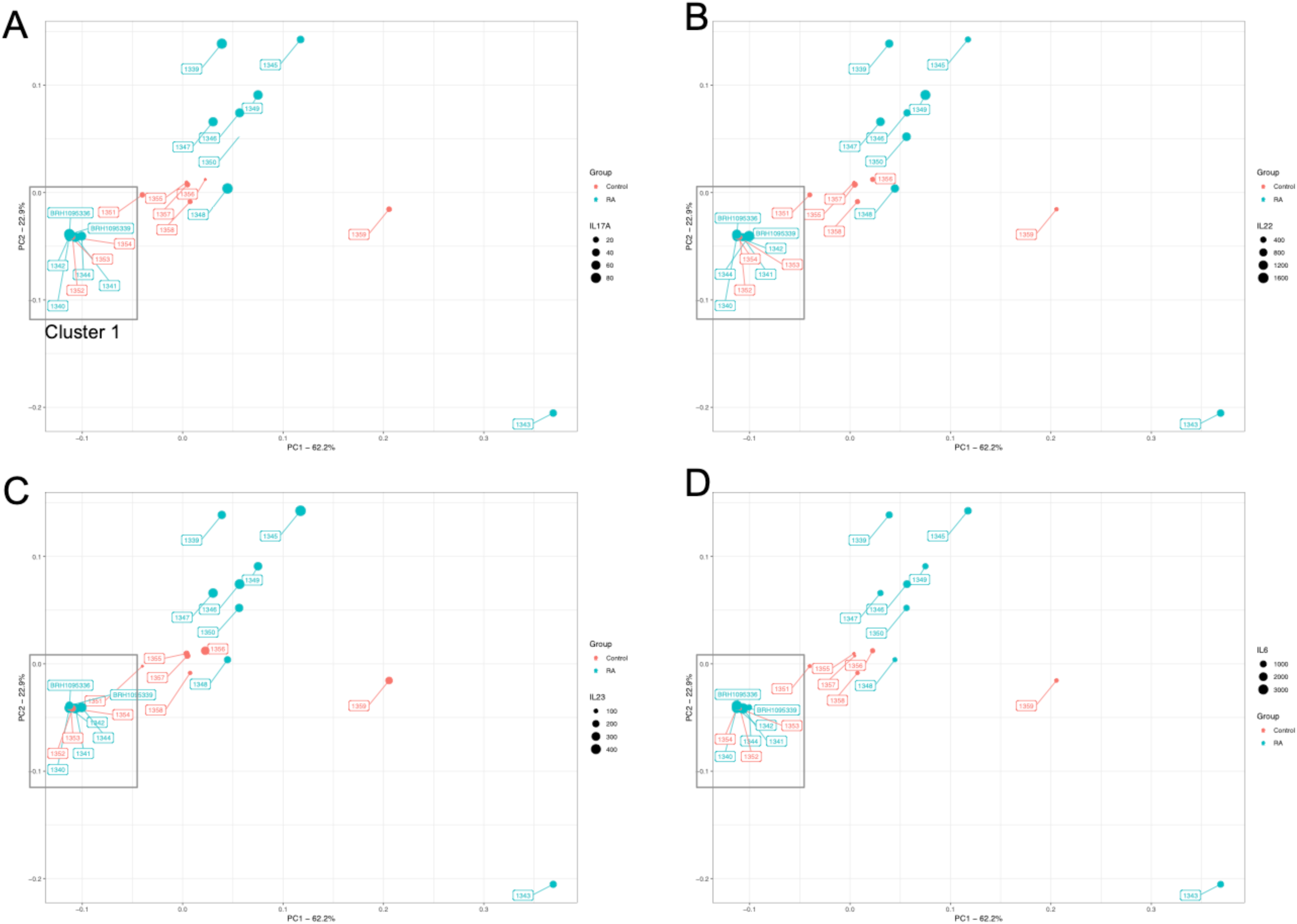
Principal Coordinate Analysis plot generated from a Bray Curtis distance matrix of synovial fluid bacterial community structure for control subjects (red) and arthritic patients (blue) as determined via amplification and sequencing of the 16S rRNA gene variable region 4 (V4). Proportions of variation explained by the principal coordinates are designated on the axes. PCoA found that the maximal variation were 62.2% (PC1), and 22.9% (PC2). The microbiome of samples that appear in close proximity to each other are more similar in composition. The data points in each pane are sized according to the synovial concentration of IL-17A (A), IL-22 (B), IL-23 (C), and IL-6 (D).

At the phylum level, the synovial fluid microbiome was predominated by *Proteobacteria* (Control = 83.5%, RA = 79.3%) and *Firmicutes* (Control = 16.1%, RA = 20.3%), and to a much lesser extent, *Actinobacteria* (Control = 0.2%, RA = 0.3%) and *Bacteroidetes* (Control = 0.1%, RA = 0.1%). At the genus level, our synovial fluid samples were predominated by the genus *Pseudomonas* (Control = 80.6%, RA = 66.1%, P = 0.30), followed by the genus *Enterococcus* (Control = 11.7%, RA = 11.4%, P = 0.95). To a lesser extent, synovial fluid samples contained members of the *Stenotrophomonas* (Control = 1.5%, RA = 4.2%, P = 0.97), *Raoultella* (Control = <0.1%, RA = 6.5%, P = 0.051) and *Bacillus* (Control = 0%, RA = 1.2%, P = 0.99) genera, the latter two being detected almost exclusively in the synovial fluid of our RA patients (**Figure 2**). Finally, unidentified taxa of the *Bacillales* order (Control = 4%, RA = 7.4%, P = 0.97) were also present in relatively high abundance yet could not be further taxonomically classified. Statistical analysis revealed that whilst the abundance of most bacterial DNA, at different taxonomic levels, was unaffected by disease status, the abundance of *Raoultella* was borderline significant (P = 0.051) and observed as the almost exclusive presence of this organism’s DNA in the synovial fluid of RA patients only (**Figure 3**).

**Figure 2.**
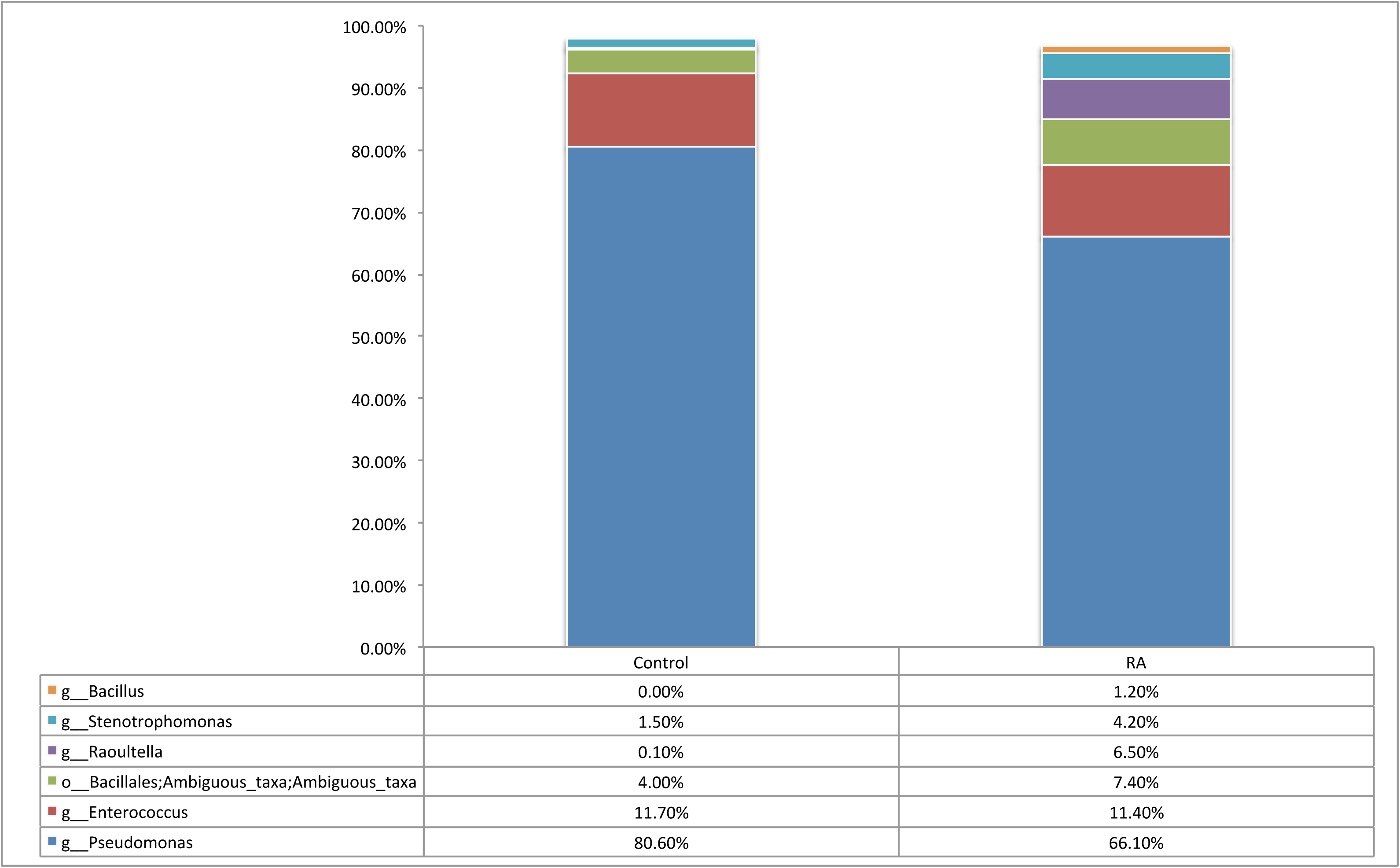
Relative abundance of genera representing >1% of the total bacterial population in synovial fluid samples of rheumatoid arthritis (RA, n = 14) and control (Control, n = 9) samples as determined by amplification and sequencing of the 16S rRNA gene variable region 4. Data are mean abundance expressed as a percentage of the total bacterial sequence count.

**Figure 3.**
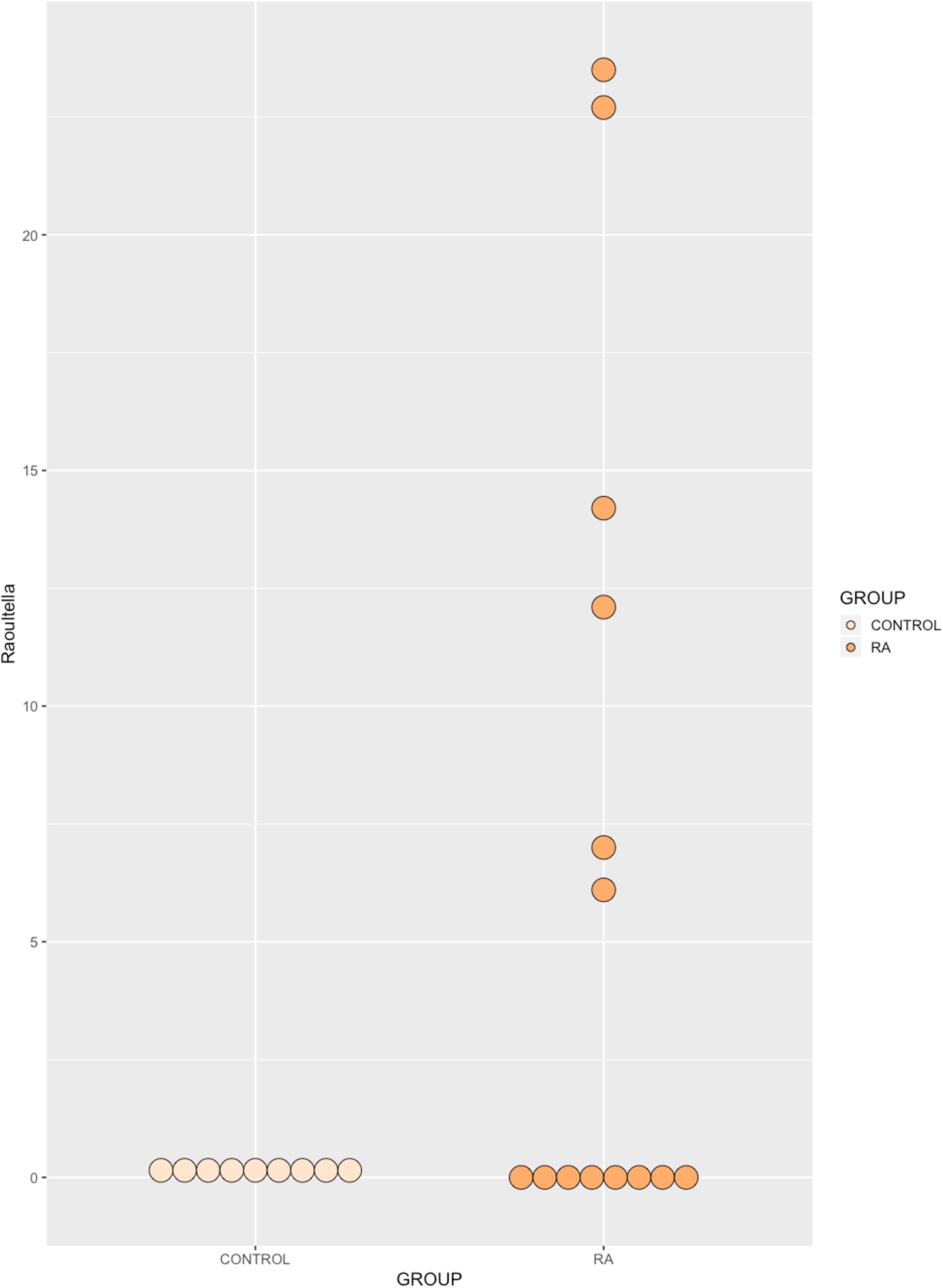
Relative abundance of the genus *Raoultella*, identified in the synovial fluid of healthy donors and RA patients by amplification and sequencing of the 16S rRNA gene variable region. A borderline significant (p = 0.051) increase in the mean abundance of *Raoultella* was identified in RA synovial fluid relative to healthy control synovial fluid. Data are individual sample abundances expressed as a percentage of the total bacterial sequence count.

### Inflammatory markers and modulated by bacterial microbiome PCoA cluster

Based upon the clustering of samples observed following PCoA (**Figure 1**), we investigated whether the samples in “cluster 1”, had significantly different inflammatory marker levels to the remaining samples, in addition to different microbial populations. Interestingly, IL-6 was significantly higher in PCoA cluster 1 compared with the remaining samples (Two Way ANOVA; P = 0.030) yet all other inflammatory markers were unaffected by PCoA cluster (P > = 0.05).

### Characterisation of fungal populations via ITS2 sequencing of synovial fluid

The presence of fungal DNA in synovial fluid was evaluated via PCR amplification and sequencing of the fungal ITS2 gene. An average of 50,000 reads were generated for each of the samples; 52,113 reads in the RA samples, and 49,929 reads in the Control samples. Although the RA samples generated more reads on average, this difference was not statistically significant (P = > 0.05). Following taxonomic classification using the Nephele platform as described above, our first approach utilised Principal Coordinates Analysis (PCoA) to reduce the complexity of the data obtained, and to visualise any obvious separation between the two experimental groups (**Figure 4**).

**Figure 4.**
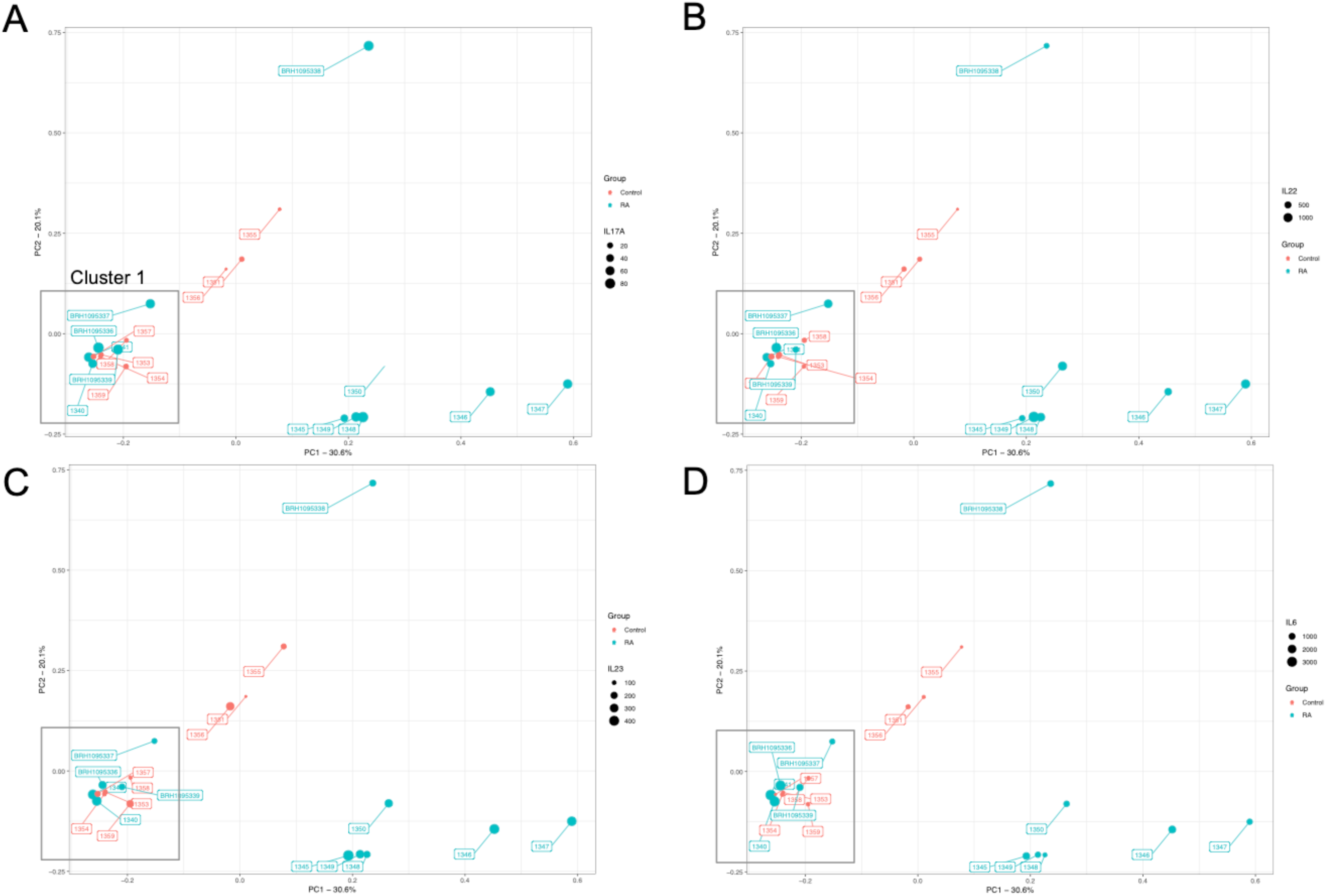
Principal Coordinate Analysis plot generated from a Bray Curtis distance matrix of synovial fluid fungal community structure for control subjects (red) and arthritic patients (blue) as determined by amplification and sequencing of the ITS2 gene. Proportions of variation explained by the principal coordinates are designated on the axes. PCoA found that the maximal variation were 30.6% (PC1), and 20.1%. The microbiome of samples that appear in close proximity to each other are more similar in composition. The data points in each pane are sized according to the synovial concentration of IL-17A (A), IL-22 (B), IL-23 (C), and IL-6 (D).

At the phylum level, synovial fluid was found to be predominated by members of the Basidiomycota (Control = 53.9%, RA = 46.9%) and Ascomycota (Control = 35.1%, RA = 50.8%) phyla. At the genus level, synovial fluid samples were predominated by the genus *Malassezia*, which accounted for 45.3% and 40.9% of the total number of fungal sequences identified in the Control and RA donors respectively (P = 0.8). To a lesser extent, synovial fluid samples also contained organisms from the genera *Penicillium* (Control = 0.7%, RA = 15%, P = 0.6), *Candida* (Control = 11.9%, RA = 0.00%, P = 0.4), *Pichia* (Control = 8.9%, RA = 0.00%, P = 0.4), *Aspergillus* (Control = 0.2%, RA = 6.2%, P = 0.14), Cladosporium (Control = 5.1%, RA = 0.1%, P = 0.019), and *Debaryomyces* (Control = 4.7%, RA = 0.4%, P = 0.7). Furthermore, genus level classification revealed the presence of specific unclassified organisms belonging to the classes *Tremellomycete*s (Control = 3%, RA = 2.8%, P = 0.98), and *Agaricomycetes* (Control = 3.2%, RA = 2.2%, P = 0.3), and the orders *Hypocreales* (Control = 0.00%, RA = 26.4%, P = 0.012) and Malasseziales (Control = 1.2%, RA = 0.1%, P = 0.002) (**Figure 5**). Statistical analysis revealed that whilst the abundance of most fungal DNA was unaffected by disease state, select organisms were differentially abundant or present (**Figure 6**). An unclassified member of the *Hypocreales* order (P = 0.012) and a member of the genus *Aspergillus* was detected almost exclusively in RA synovial fluid samples only (P = 0.14 including all data, P = 0.057 excluding samples where <1% of reads mapped to this taxa). Conversely, a member of the order *Malasseziale*s (P = 0.002), and the genus *Cladosporium* (P = 0.019) were observed almost exclusively in control synovial fluid.

**Figure 5.**
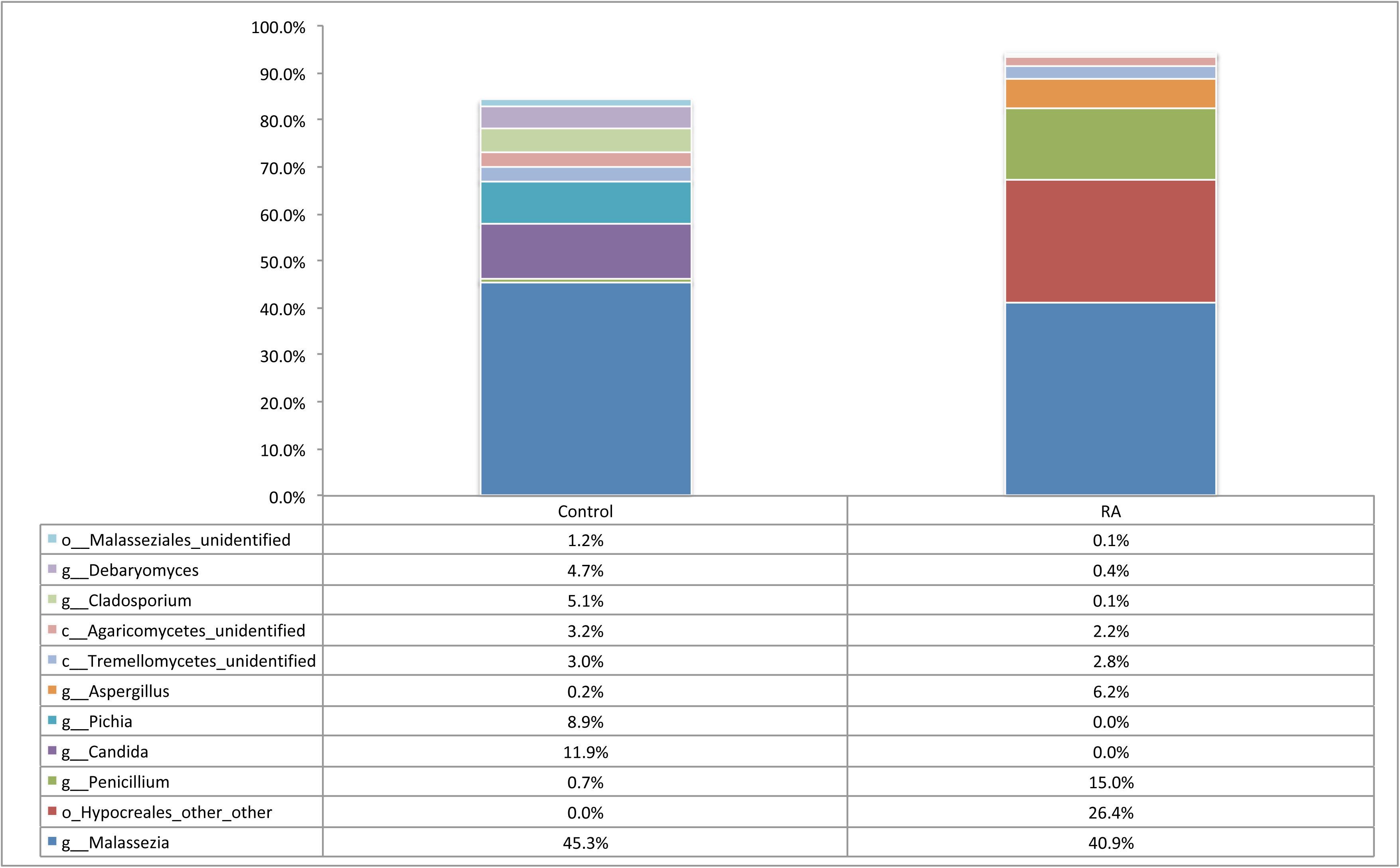
Relative abundance of taxa representing >1% of the total fungal population in each synovial fluid sample (RA, n = 12) and (Control, n = 9) as determined by amplification and sequencing of the fungal ITS2 gene. Data are mean abundance expressed as a percentage of the total fungal sequence count.

**Figure 6.**
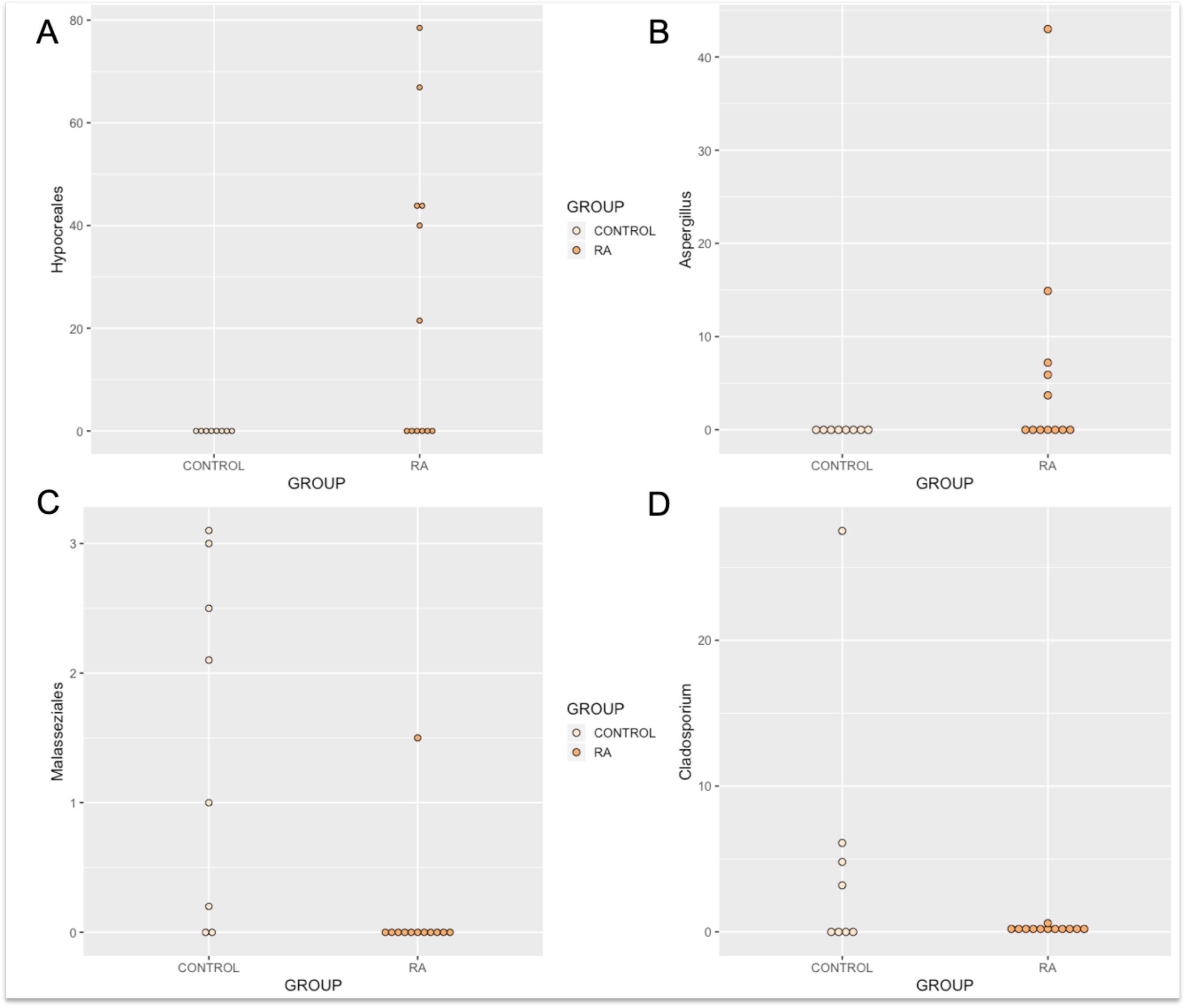
Relative abundance of (A) *Hypocreales*, (B), *Aspergillus*, (C) *Malasseziales*, and (D) Cladosporium identified in the synovial fluid of RA patients and healthy control subjects. Data determined by the amplification and sequencing of the ITS2 gene. (A) The relative abundance of order *Hypocreales* was increased in synovial fluid of RA patients relative to healthy control synovial fluid. (B) The relative abundance of genus Aspergillus was increased in RA synovial fluid samples compared with healthy subjects (borderline significant). (C), and (D) The abundance of the order *Malasseziales* and genus *Cladosporium* were significantly decreased in the synovial fluid of RA patients in comparison to the synovial fluid of healthy control subjects. Data are individual expression values expressed as a percentage of the total fungal read count in each sample. The statistical significance between groups was determined by unpaired t test where P□≤□0.05.

### Inflammatory markers and fungal microbiome PCoA cluster

Based solely upon the clustering of samples observed following PCoA, we investigated whether samples in PCoA “cluster 1” had significantly different inflammatory marker profiles, in addition to different microbial populations, than the remaining samples. IL-6 was significantly higher in PCoA cluster 1 compared with the remaining samples (Two Way ANOVA; P = 0.016). Further, the interaction between PCoA cluster and disease status (control or RA) was also significant (P = 0.048). All other inflammatory markers were unaffected by PCoA cluster (P > = 0.05).

## Discussion

Mounting evidence suggests pan-microbiome dysbiosis in patients with RA. Moreover, various studies spanning many decades link the development of RA with infections of the oral tissues or urinary system, and report the presence of microorganisms or microorganism-derived genetic material in the synovial fluid; a compartment classically considered to be sterile. To our surprise, only one study has investigated the potential for a “synovial fluid microbiome” using next generation sequencing [5]. To this end, we investigated the presence of bacterial and fungal DNA in the synovial fluid of rheumatoid arthritis (RA) patients and healthy control subjects through amplification and sequencing of the 16S rRNA and ITS2 genes.

### Evidence of bacteria

Synovial fluid samples from 25 subjects were analysed in this study and our analyses revealed the presence of a complex synovial microbiome community in health and disease. Bacterial 16S rRNA was detected in fourteen out of sixteen (87.5%) RA patients, and nine out of nine (100%) healthy control subjects. At the phylum level, the synovial fluid microbiome was predominated by Proteobacteria (81.1%) and Firmicutes (18.5%), and to a much lesser extent, Actinobacteria (0.3%) and Bacteroidetes (0.1%). These phyla were also identified as the most abundant in the only other comparable study [5], albeit in slightly different proportions. Whilst the major phyla identified appear similar to those of the blood microbiome [59], they are present in the synovial fluid at different levels, suggesting a distinct community structure.

At the genus level, our synovial fluid samples were predominated by DNA of the genus *Pseudomonas* and *Enterococcus*, to a lesser extent, unidentified taxa of the *Bacillales* order and *Stenotrophomonas* genus. The genera *Raoultella* and *Bacillus* were detected almost exclusively in the synovial fluid of our RA patients (**Figure 2 and 3**). DNA of *Pseudomonas* species, which was the most common genus to be identified in the study cohort, has previously been identified in the synovial fluid of patients with RA [42, 60].

Whilst the bacterial communities characterised in the disease and control states were largely consistent, the presence of the genus *Raoultella* was largely restricted to the RA synovial fluid in our study (present only in 4 of our control samples with read counts < 0.3% yet detected in 6 of our RA patient samples with read abundances in the range 6.1 – 23.5%). Interestingly, when present, *Raoultella* often occupied a large proportion of the total bacterial read count. This is the first study to describe the presence of DNA from the genus *Raoultella* in RA synovial fluid samples, however, two major species are capable of causing infections in man: *Raoultella ornithinolytica* and *Raoultella planticola*, and have been shown to be capable of reaching the articular joint compartment [61–63]. Moreover, *Raoultella* are characterised as histamine generating bacteria [64], a key pro-inflammatory molecule found in RA blood, articular cartilage, synovial fluid and synovial tissue [65]. It is plausible that the presence and or indeed abundance of *Raoultella* DNA may correlate with specific disease features (for example; disease stage, severity, current disease activity, treatment regimen), and we therefore plan to explore this relationship further in the future, particularly with the aim of identifying the origin of the DNA found.

With regards the numerous reports detailing a link between oral and urinary tract associated bacteria and RA, we reviewed the results of our taxonomic classification looking specifically for the presence and abundance of these organisms. Bacteria of the genus *Porphyromonas* were largely absent with the exception of three synovial fluid samples (Control = 2, RA =1), which returned a small number of reads mapping to this genus. These results differ from those of Zhao and colleagues who report the presence of *Porphyromonas* in all of the samples they sequenced [5], irrespective of disease state [5]. No reads mapping to the genus *Proteus* were identified in any sample analysed, however numerous other members of the order Enterobacteriales to which *Proteus* belongs were identified, including *Raoultella*. It is important to note here that none of our RA patients reported oral or urinary conditions, and it is therefore plausible that the above bacteria are present only during active oral or urinary disease. A separate study is therefore warranted to further investigate these phenomena including RA patients (without co-morbidities), RA patients with poor oral health, and patients with periodontitis alone (without RA).

### Evidence of fungi

Fungal DNA was identified in 12 out of 16 (75%) RA samples, and 8 out of 9 (88.8%) healthy controls. Our various experimental controls consistently failed to produce a visible band following PCR and agarose gel electrophoresis. Furthermore, DNA quantification using the Qubit 3.0 high-sensitivity DNA kit (Invitrogen) was consistently below the limit of detection. These indicate that the fungal DNA identified originates from within the samples rather than from contaminants introduced. At the phylum level, synovial fluid was predominated by members of the Basidiomycota and Ascomycota phyla. At the genus level, synovial fluid samples were predominated by the genus *Malassezia*, which accounted for 45.3% and 40.9% of the total number of fungal sequences identified in the Control and RA donors respectively. To a lesser extent, synovial fluid samples also contained organisms from the genera Penicillium, *Candida, Pichia, Aspergillus*, Cladosporium, and *Debaryomyces*. Furthermore, genus level classification revealed the presence of specific unclassified organisms belonging to the classes *Tremellomycete*s and *Agaricomycetes*, and the orders *Hypocreales* and Malasseziales (**Figure 6**).

Again we identified key taxa that appeared to be differentially present or abundant dependent on disease status. The order *Hypocreales* was selectively abundant in RA patients (7 / 12) and absent from healthy control synovial fluid. Interestingly, when present, members of the Hypocreales order occupied a large proportion of the total fungal read count (~ 40%). Evidence for the presence of members of this order have been previously identified in the healthy human oral cavity [66] and found to be ubiquitous in human plasma [67], however no reports directly link this order with RA.

Further, the abundance of genus *Aspergillus* was borderline significant; detected more commonly in the synovial fluid of RA patients, and occupying an average of ~ 10.0% of the read population when present. *Aspergillus* has been identified in the healthy human gut [68] and oral cavity [66] in various studies, and is a prevalent opportunistic pathogen of the lungs, particularly affecting immunocompromised individuals, where it causes a range of pulmonary diseases [69]. Furthermore, *Aspergillus* galactomannan antigen level has been shown to elevate in a non-specific manner among patients with RA [70], further hinting towards a possible association.

The presence members of the genus Cladosporium were largely absent in the synovial fluid of all RA patients in comparison to the synovial fluid of healthy control subjects (present in 5 / 8 healthy control patients in comparison with 2 / 12 RA samples). Cladosporium has been shown to colonise the healthy human gut [68] and oral cavity [66], and is encountered commonly in human clinical samples [71], thus evidence of Cladosporium DNA in the healthy patient is perhaps not unexpected. That said, it’s absence from the synovial fluid of the majority of RA patients may suggest an altered fungal community in some distant microbial niche, that presents as a reduction in translocated Cladosporium DNA. These findings all warrant further investigation.

*In-vivo* studies have demonstrated that mycotoxins may play a role in the pathogenesis of RA [72]. Furthermore, cells such as Th17 that are involved in the immunity against fungi are also implicated in the pathogenesis of RA [73]. These indicate that fungal presence in synovial fluid of patients with RA may contribute to pathogenesis. However, as fungal DNA was also found in healthy controls, further studies are needed, with quantification of the load of fungal DNA and taxa specific components.

### Changes in microbiome with synovial fluid cytokine levels

Numerous cytokines have been implicated in the pathogenesis of RA. Levels of IL-17-A, IL-22, IL23, and IL-6 have been identified to be higher in diseased joints than in healthy [74, 75], and IL-6 has been identified to play a key role in the pathogenesis of RA. It is implicated in pathogenesis of local joint disease as well as systemic manifestations [76, 77]. Serum and synovial fluid IL-6 levels correlate with disease severity and activity [78, 79]. We found that IL-6 levels were higher among a sub-cluster of the RA population defined according to the fungal and bacterial microbiome. As mentioned earlier, Th17 cells are important in the immunity against fungal pathogens, and IL-6 is involved in the differentiation of lymphocytes to Th17 cells [77]. This could indicate that the differences in the fungal microbiome within patients with RA might be related to disease severity through an IL-6 – Th17 mediated pathway.

### Limitations

Bacterial DNA was identified in 14/16 patients with RA, and in all disease free controls whereas fungal DNA was identified in 12 out of 16 RA samples, and 8 out of 9 healthy controls. These results are consistent with other published works that have previously reported the presence of bacterial DNA in the majority, but not necessarily all, samples investigated [80, 81]. A potential limitation of this study is that we did not elect to sequence of experimental controls, and relied on the absence of a detectable signal following agarose gel electrophoresis and QuBit analysis using the high sensitivity DNA kit. Whilst is plausible that sequencing could have revealed the presence of low abundance target DNA introduced during our experimental procedures we suggest that this (potential) DNA would be (1) present at a much lower concentration (given our experimental samples were clearly visible following gel electrophoresis relative to the absence of any bands for our experimental controls), and (2) consistent given that all samples were processed in parallel. Whilst we acknowledge this potential limitation, it does not detract from our finding of statistically different levels of specific bacteria between the two experimental groups, despite all samples being processed at the same time utilising the same specific batches of reagents.

## Conclusions

This is one of the first studies to report the existence of a synovial fluid microbiome, and to determine that this is modulated by disease status (RA) as are other classical microbiome niches. Our recent work on the blood microbiome [4], and that of others [82–92], characterised a complex community of organisms (or the genetic material thereof, more specifically) that likely originate in one of the classical microbiome niches (the gut, mouth, urogenital tract, skin) and reach the circulation through a process termed atopobiosis. We hypothesise that the same process may be true of the community we characterised here in the synovial fluid, with the circulatory system acting as a conduit between their usual place of habitation and the synovial compartment. Further studies incorporating pan-microbiome analysis are required to support these data.

### Preprint

This manuscript has been published as a preprint bioRxiv [93].

